# A Leaky Human Colon Model Reveals Uncoupled Apical/Basal Cytotoxicity in Early *Clostridioides difficile* Toxin Exposure

**DOI:** 10.1101/2022.10.13.511617

**Authors:** Meryem T. Ok, Jintong Liu, R. Jarrett Bliton, Caroline M. Hinesley, Ekaterina Ellyce T. San Pedro, Keith A. Breau, Ismael Gomez-Martinez, Joseph Burclaff, Scott T. Magness

## Abstract

**Background & Aims:** *Clostridioides difficile* (*C. difficile*) toxins A (TcdA) and B (TcdB) cause antibiotic-associated colitis and increase morbidity and mortality. Accurate in vitro models are necessary to detect early toxicity kinetics, investigate disease etiology, and develop pre-clinical models for new therapies. Properties of cancer cell lines and 3D organoids inherently limit these efforts. Here, we develop adult stem cell-derived monolayers of differentiated human colonic epithelium (hCE) with barrier function, investigate the impact of toxin application to apical/basal aspects of monolayers, and evaluate whether a leaky epithelial barrier enhances toxicity.

**Methods:** Single-cell RNA-sequencing (scRNAseq) mapped *C. difficile*-relevant genes to cell lineages across the human gut. Transcriptomics informed timing of stem cell differentiation to achieve in vitro colonocyte maturation like that observed in vivo. Transepithelial electrical resistance (TEER) and fluorescent dextran permeability assays measured cytotoxicity as barrier loss post-toxin exposure. Leaky epithelial barriers were induced with diclofenac.

**Results:** scRNAseq demonstrated broad and variable toxin receptor expression across the human gut lineages. Absorptive colonocytes displayed generally enhanced toxin receptor, Rho GTPase, and cell junction expression. 21-day differentiated Caco-2 cells remained immature whereas hCE monolayers were similar to mature colonocytes. hCE monolayers exhibited high barrier function after 1-day differentiation. Basal TcdA/B application to monolayers caused more toxicity and apoptosis than apical exposure. Diclofenac induced leaky hCE monolayers and enhanced toxicity of apical TcdB exposure.

**Conclusions:** Apical/basal toxicities are uncoupled with more rapid onset and increased magnitude of basal toxicity. Leaky paracellular junctions enhance toxicity of apical TcdB exposure. hCE monolayers represent a physiologically relevant and sensitive culture system to evaluate the impact of microbial toxins on gut epithelium.

## INTRODUCTION

*Clostridioides difficile* is a toxin-producing bacterium that causes *C. difficile* infection (CDI), with symptoms ranging from mild diarrhea to severe colitis and associated with life-threatening illnesses such as toxic megacolon and bowel perforation.^1^ CDI has a tremendous economic burden, costing an estimated $6.3 billion in annual healthcare expenses in the U.S. alone.^2^ Patients with greater CDI risk include those who are immunocompromised, receiving broad-spectrum antibiotics, or battling a leaky gut, often due to underlying chronic illness such as inflammatory bowel disease (IBD).^2,3^ While fidaxomicin and vancomycin remain first-line antibiotics for patients with CDI, the high treatment costs,^4^ emergence of antibiotic-resistant strains,^5^ and increase in recurrence rates ^6^ especially for hypervirulent strains (≥ 23%)^6,7^ are growing problems.

*C. difficile* damages the colonic epithelium by secreting Toxin A (TcdA), Toxin B (TcdB), and binary toxin *C. difficile* transferase (CDT). CDT is produced by approximately 20% of *C. difficile* strains, but it receives less clinical attention given that CDT-producing strains lacking TcdA and TcdB are nontoxigenic in vitro.^8,9^ TcdA and TcdB (TcdA/B) are homologous glucosyltransferases that bind to various host cell receptors, enter cells via endocytosis, and inactivate Rho-family GTPases by monoglucosylation.^10^ This ultimately results in cytoskeletal disassembly, tight junction collapse, cytopathic cell rounding, and eventual apoptosis.^11–15^ While the TcdA/B receptor distribution likely impacts the onset of cytopathic events, this distribution has not been thoroughly mapped across the proximal-distal axis of the human gut or across epithelial cell lineages. A comprehensive map of *C. difficile* toxin receptors across all undifferentiated and differentiated lineages would address a major gap in knowledge and serve as a foundation to better understand mechanisms of toxin interactions with the epithelium, disease onset, progression, and resolution.

Alternatives to antibiotic therapies are becoming an attractive approach to treat CDI; however, physiologically relevant preclinical models of the human colon to test new therapies are lacking. Historically, *C. difficile* toxicity models relied on non-intestinal cell types (e.g., Vero, HeLa) or colon cancer cell lines (e.g., T84, HT-29, Caco-2).^16–19^ However, colon cancer cells do not express physiological transporters/carriers found in healthy human colonic epithelium (hCE), and they phenotypically resemble small intestinal enterocytes.^20,21^ While TcdB is more cytotoxic than TcdA in T84 cells and non-intestinal epithelium,^22,23^ TcdA is more potent than TcdB in 3D human intestinal organoids (HIOs) and jejunal enteroid-derived monolayers.^24,25^ Although a significant improvement in physiological accuracy, 3D organoids—in which the apical cell surface faces inward inside an enclosed epithelium—do not allow easy simultaneous access to apical/basal cell surfaces to apply toxins or assess barrier function using transepithelial electrical resistance (TEER) or permeability assays. Moreover, HIOs are induced to differentiate from pluripotent stem cells in vitro and the resulting organoids are more similar to fetal small intestine^26^ than colon, which is the predominant site of CDI,^27^ giving these findings limited generalizability to the sensitivity of adult differentiated hCE to TcdA/B. Thus, more physiologically accurate culture systems are essential to mimic in vivo colonic epithelium and investigate the impact of TcdA/B on hCE.

Currently, there are no established quantitative culture systems that simultaneously evaluate the apical versus basal impact of TcdA/B on hCE. *C. difficile* toxins are secreted in the lumen and thought to interact first with the apical aspect of the intestinal or colonic epithelial barrier.^28^ However, receptor studies in colon cancer cell lines^28,29^ and in non-human, non-intestinal epithelial cells^30^ reveal that some TcdA receptors locate to the basal side of the cell, making them inaccessible to TcdA when there is an intact physiological epithelial barrier. This raises the possibility that a healthy epithelial barrier may be more refractory to apical TcdA cytotoxicity than previously believed, but this has yet to be tested in hCE. Likewise, while some TcdB receptors localize to the basal aspect of the epithelium,^31–33^ their full apical/basal distribution remains unclear,^34,35^ though functional studies in HIOs show distinct apical and basal differences in barrier disruption.^24^ Thus, we designed a model to quantify the magnitude and time-course of apical/basal toxicity, which could inform whether early clinical interventions should target apical versus basal hCE.

Here, we establish a physiologically relevant in vitro model of *C. difficile* TcdA/B toxicity using primary adult intestinal stem cell (ISC)-derived hCE monolayers. Building upon our previous methods of culturing primary ISCs from healthy human donor organs and differentiating these ISCs into a functional epithelial layer,^36–38^ we develop a highly sensitive assay system for evaluating early-stage intestinal barrier injury and late-stage cytotoxic events associated with TcdA/B. Our tissue engineering approach focuses on producing an epithelium that closely mimics in vivo colonic tissue, which we use to assess *C. difficile*-relevant transcriptional and phenotypic differences between hCE, human small intestinal epithelium (hSIE), and Caco-2 cells. Using this system, we test the temporal dynamics of apical and basal TcdA/B toxicity, measure the effect of a leaky epithelium on toxin activity, and present biological insights into the impact of apical and basal cytotoxicity of TcdA/B.

## RESULTS

### A Single-Cell Survey of Human Gut Epithelium Reveals Organ and Lineage Variability of *C. difficile* Toxin-Relevant Genes

Expression of *C. difficile*-relevant gene sets have not been characterized across the epithelial lineages of the small intestine and colon. To address this, we interrogated single-cell transcriptomic data across all lineages in small intestine and colon from three healthy human adult donors **(Fig. 1A)**^39^ and evaluated the magnitude of toxin receptor expression **(Fig. 1B)**.^28, 29, 31, 34, 40, 41^ In general, the ISC, transit-amplifying (TA), and absorptive lineages in the small intestine and colon exhibited more toxin receptor expression and with higher magnitudes compared to the secretory lineages. Notably, follicle-associated epithelium (FAE) uniquely expressed high levels of the TcdB receptor *FZD7* **(Fig. 1B)**. Lineage-specific *C. difficile* toxicities have not been evaluated; thus, these findings provide a foundation to investigate such questions.

**Figure 1.**
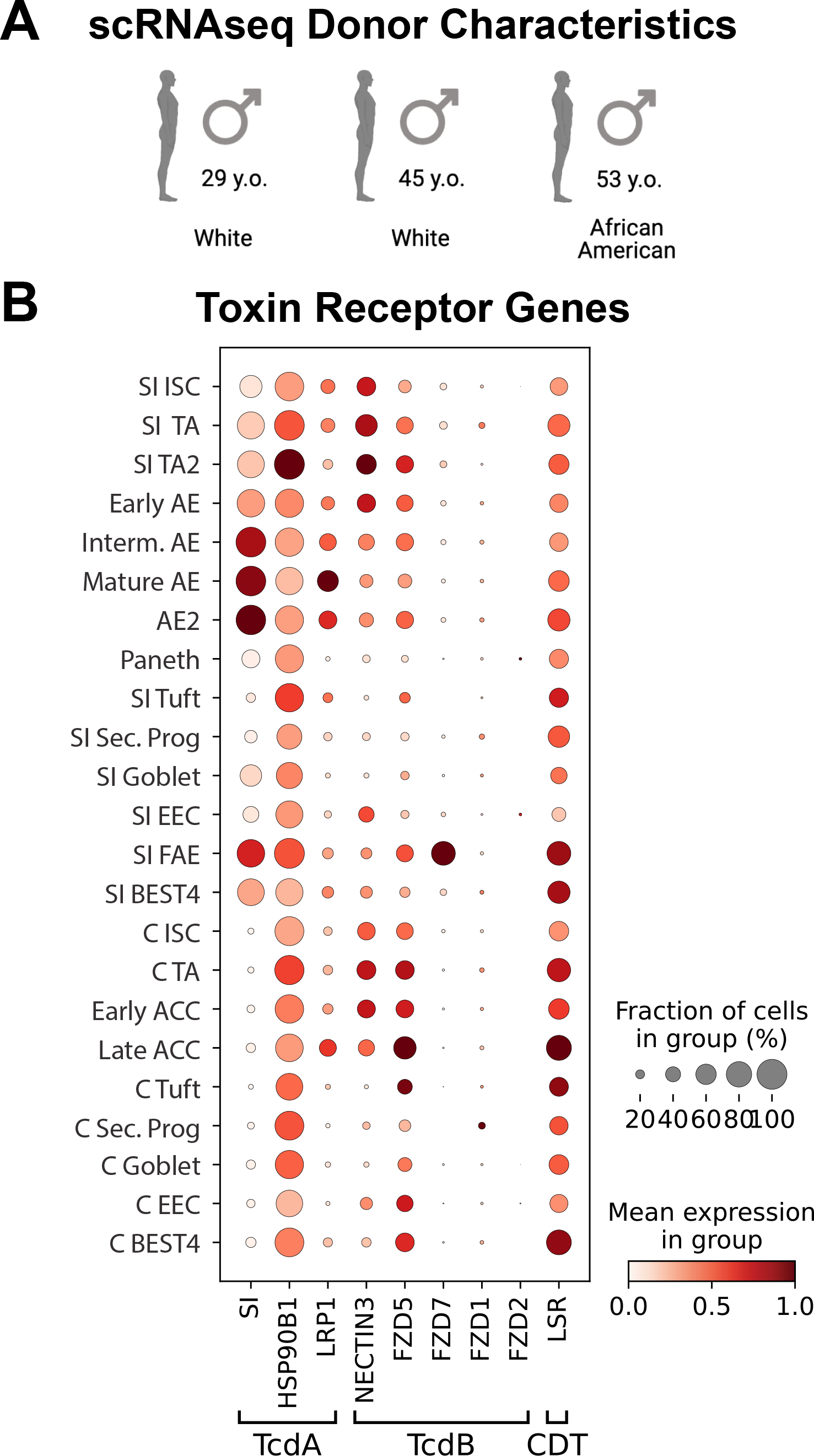
Single-cell transcriptomics demonstrates variations in expression patterns and magnitude of *C. difficile* toxin receptor genes across all intestinal epithelial cell lineages. **(A)** Healthy human organ donor demographic information for single-cell RNAseq data. **(B)** Dotplot showing the fraction of expressing cells (circle size) and magnitude (color) of expression for known *C. difficile* toxin receptor genes across all small intestinal and colonic epithelial cell lineages. SI, Small Intestine; ISC, Intestinal Stem Cell; TA, Transit-Amplifying; AE, Absorptive Enterocyte; Interm., Intermediate; Sec. Prog, Secretory Progenitor; EEC, Enteroendocrine Cell; FAE, Follicle-Associated Epithelium; C, Colon; ACC, Absorptive Colonocyte.

Next, we sought to evaluate the *C. difficile*-relevant genes in the two key cell types involved in the initiation and resolution of *C. difficile* infections. The absorptive lineages are the predominant cell types in the small intestine and colon and experience first-line exposure to *C. difficile*, and the ISCs are responsible for regenerating the epithelium after damage. We surveyed curated gene sets for toxin receptor, Rho GTPase, and tight junction expression in ISCs and mature absorptive cells in six regions across the proximal-distal axis of the small intestine (Duodenum, Jejunum, Ileum) and colon (Ascending, Transverse, Descending) **(Fig. 2A)**. For the toxin receptor genes, sucrase-isomaltase is a small intestine-specific brush border enzyme;^42^ thus, it is not expected to be meaningfully expressed in the colon, as demonstrated by the data **(Fig. 2A)**. When comparing the remaining receptors between small intestinal and colonic ISCs, there was no difference in *HSP90B1* and *LSR*, *FZD5* was significantly higher in colonic ISCs, and *LRP1* and *NECTIN3* were higher in small intestinal ISCs **(Fig. 2B)**. In mature absorptive cells of the small intestine and colon, all receptors except for *LRP1* and small intestinal-specific *SI* were significantly higher in colon **(Fig. 2B)**. For classic Rho-family GTPases (*CDC42, RAC1*, and *RHOA*),^43–45^ which are terminal targets of *C. difficile* toxins, no specific gene demonstrated statistically significant differences between small intestinal and colonic ISCs **(Fig. 2C, D)**. However, in mature absorptive lineages, *RAC1* was modestly but significantly higher in small intestine, and *CDC42* and *RHOA* were higher in the colon **(Fig. 2C, D)**.

**Figure 2.**
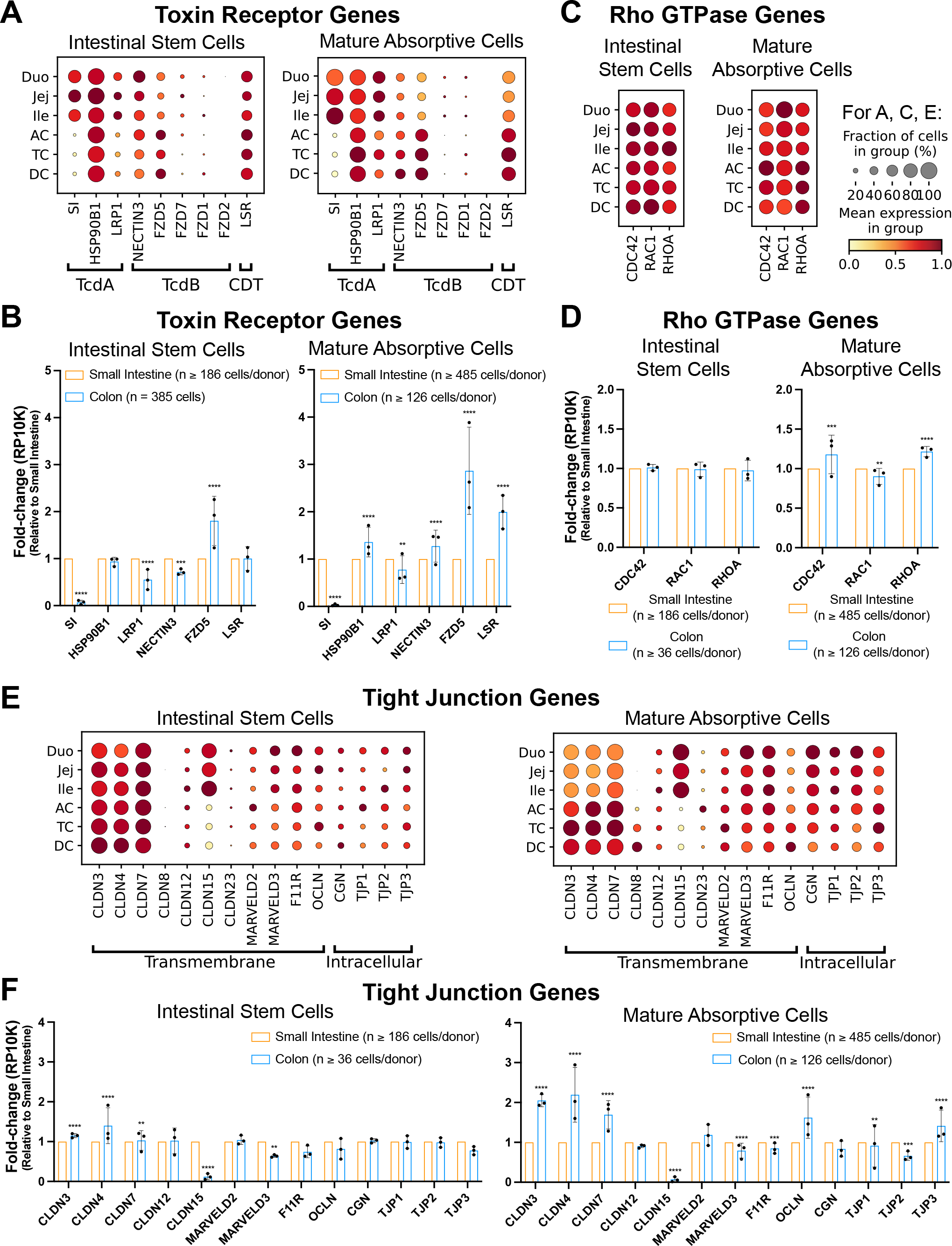
Single-cell RNAseq survey of *C. difficile*-toxin relevant genes reveals organ and lineage variability. **(A)** Dotplots comparing *C. difficile* toxin receptor genes across small intestinal and colonic epithelial regions in stem cells (left) and mature absorptive cells (right) and **(B)** corresponding significance bar graphs. **(C)** Dotplots comparing Rho GTPase family genes by intestinal region in stem cells (left) and mature absorptive cells (right) and **(D)** corresponding significance bar graphs. **(E)** Dotplots comparing curated tight junction genes by intestinal region in stem cells (left) and mature absorptive cells (right) and **(F)** corresponding significance bar graphs. All bar graphs show fold-change in reads per 10,000 genes (RP10K) across 3 donors for colon normalized to small intestine, mean ± SD. \**q* < 0.05, \*\**q* < 0.01, \*\*\**q* < 0.001, \*\*\*\**q* < 0.0001. Duo, Duodenum; Jej, Jejunum; Ile, Ileum; AC, Ascending Colon; TC, Transverse Colon; DC, Descending Colon.

We analyzed tight junction proteins, which are known to be disrupted as a terminal effect of toxin-induced Rho GTPase glucosylation.^14^ A subset of 34 genes were selected from a published curated gene set based on classification of ‘tight junction’ or ‘tight junction-associated’ genes.^46^ Fifteen genes with the highest mean expression in mature small intestinal and colonic absorptive lineages were grouped by transmembrane versus intracellular proteins and their expression was characterized across regions and lineages of the small intestine and colon **(Fig. 2E)**. Claudin (*CLDN*) genes *CLDN3*, *CLDN4*, and *CLDN7*, which aid in forming selective barriers to macromolecules (such as *C. difficile* toxins) and ions,^47^ are highly expressed across all ISCs and mature absorptive cells, with notably higher expression in mature absorptive lineages of the colon **(Fig. 2F)**. *CLDN15* showed unique and high expression in the ISCs and mature absorptive lineages of the small intestine. ISCs and mature absorptive lineages across all small intestinal and colonic regions demonstrated broad expression of the remaining tight junction genes, which were generally expressed at lower magnitude than the claudins **(Fig. 2F)**. These findings highlight variability in small intestinal and colonic regions and lineages that could lead to functional differences in intestinal epithelial barrier function and likely influence CDI onset and progression.

### Transcriptomics Show hCE Monolayers Mature Over Time to Mimic In Vivo Mature Absorptive Colonocytes

With the single-cell transcriptomic data serving as the in vivo benchmark for physiological relevance, we sought to determine whether colonic ISCs could be differentiated in vitro to closely mimic the absorptive colonocytes in vivo. We also compared the transcriptomics of differentiated hCE monolayers to differentiated Caco-2 monolayers in an effort to establish which model more closely aligned to in vivo transcriptomic lineage states. To differentiate hCE monolayers, primary crypt-derived ISCs from the descending colon (DC) were first expanded on collagen-coated hydrogels as previously described.^38^ DC tissue was chosen given that CDI primarily manifests as colitis in the distal colon.^27^ Once confluent, these cells were then plated onto Matrigel-coated Transwell™ inserts in expansion media (EM) containing ISC growth factors (Wnt3a, R-spondin 3, and Noggin) for 4 days during which they continued to proliferate to form a confluent monolayer. To differentiate the colonic ISCs, differentiation media (DM) was applied to ISCs for an additional 4 days **(Fig. 3A)**. RNA was isolated from hCE monolayers at various timepoints (EM Day 1, EM Day 4 = DM Day 0, DM Day 2, and DM Day 4), and transcriptomics was used to verify differentiation and characterize cell maturation.

**Figure 3.**
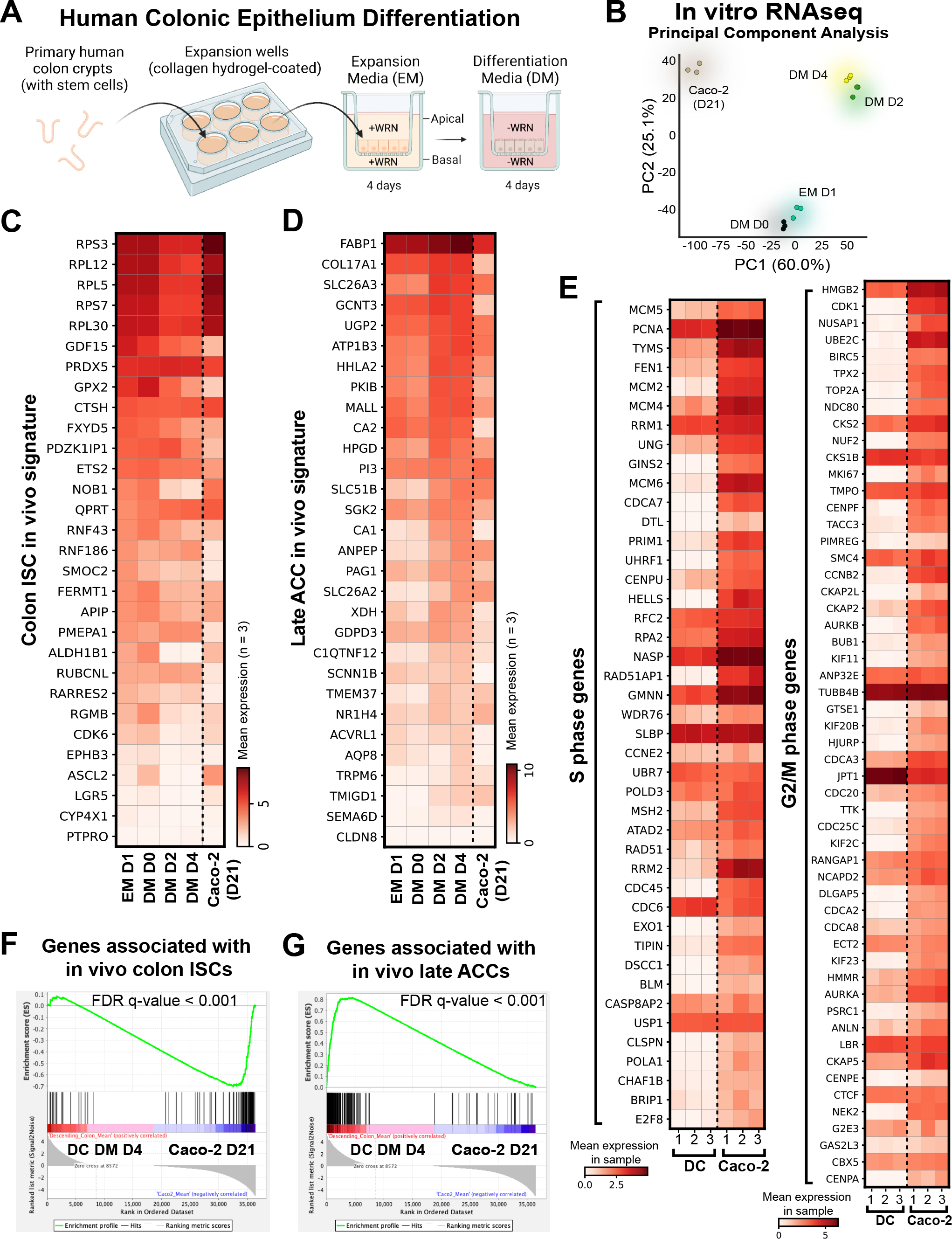
Bulk RNAseq reveals Caco-2 monolayers remain stem-like and hCE monolayers mature over time to mimic in vivo late absorptive colonocytes. **(A)** Schematic of expansion and differentiation of human colonic epithelial (hCE) monolayers on Transwells. **(B)** Principal component analysis of sequenced transcriptomes. Matrix plots comparing hCE and Caco-2 cells (n = 3 Transwells each) using top 30 differentially expressed genes (DEGs) for **(C)** colon intestinal stem cells (ISCs) and **(D)** late absorptive colonocytes (ACCs) versus the rest of the colon, and **(E)** S-phase and G2/M-phase genes in descending colon (DC) vs. Caco-2 cell monolayers. Gene set enrichment analysis for DC in differentiation media (DM) Day 4 (red) and differentiated Caco-2 cells (blue) using gene sets consisting of **(F)** all in vivo colon ISC DEGs, and **(G)** all in vivo late ACC DEGs.

Principal component analysis showed subtle transcriptomic changes while in EM (EM D1, DM D0) but substantial changes along principal component (PC) 1 and PC2 after 2 days of differentiation (DM D2), indicating large expression changes between Day 0 and Day 2 of colonic ISC differentiation **(Fig. 3B)**. There were fewer transcriptomic changes between Day 2 and Day 4 of differentiation, indicating relatively stable transcriptomes during this time window. We compared hCE transcriptomics to 21-day differentiated Caco-2 to determine if Caco-2 transcriptomics tracked with the differentiated hCEs. Caco-2 cells were markedly separated from colonic ISCs and all stages of hCE differentiation in PC1, highlighting that differentiated Caco-2 monolayers are significantly different than the primary tissue-derived hCE monolayers **(Fig. 3B)**.

We further characterized the differentiation of hCE monolayers at a more granular level to determine if time of differentiation could “mature” the monolayers to a transcriptomic state that resembles mature absorptive colonocytes in vivo. We curated two gene sets from the in vivo single-cell transcriptomic data as the standard for comparison, one from the colonic ISCs and one from the mature absorptive colonocytes (i.e. late ACCs).^39^ The hypothesis was that replacement of EM with DM would shift the transcriptomic state of the monolayers away from the ISC signature and closer to the mature absorptive colonocyte signature. The data show that hCE monolayers at EM Day 1 are more closely aligned with the in vivo colonic ISC signature, but these similarities decrease over four days of differentiation where the magnitude of most colonic ISC genes is reduced **(Fig. 3C)**. By contrast, 21-day differentiated Caco-2 cells still exhibit higher expression of many colonic ISC genes, notably *LGR5* and *ASCL2* **(Fig. 3C)**.

Conversely, when gene signatures of hCE and Caco-2 monolayers are compared to the in vivo late ACC signature, hCE cells show increased magnitude of gene expression at DM Day 2 which stabilized through DM Day 4 **(Fig. 3D)**. In most instances, the 21-day differentiated Caco-2 genes more closely aligned with the expression patterns of EM Day 1 and DM Day 0, which is reflective of a persistent undifferentiated state and further highlights the less mature state of differentiated Caco-2 **(Fig. 3D)**. To characterize the extent of the undifferentiated state between differentiated hCE and Caco-2 monolayers, curated gene sets associated with S- and G2/M-phase cell cycle stages were analyzed.^48^ We hypothesized that differentiated hCE monolayers would express relatively lower levels of S- and G2/M-phase genes while Caco-2 monolayers would show upregulated expression of these genes. The data show that 21-day differentiated Caco-2 monolayers express higher levels of nearly all genes associated with cell cycling, notably the cell proliferation markers *MKI67* and *PCNA* **(Fig. 3E)**. Finally, gene set enrichment analyses were performed using the full gene signature of in vivo colon ISCs (108 genes) and in vivo late ACCs (215 genes).^39^ Consistent with our earlier observations, Caco-2 monolayers are significantly enriched for stem-like genes compared to hCE monolayers **(Fig. 3F)**, whereas hCE monolayers are significantly enriched for late ACC genes compared to Caco-2 monolayers **(Fig. 3G)**. These findings emphasize that hCE monolayers are more terminally differentiated and more aligned with the in vivo late ACC gene signature, which likely translates into better functional equivalence compared to Caco-2 cells.

### Differentiated hCE Monolayers Exhibit Stronger TEER Barrier Properties Than Caco-2 or hSIE Monolayers

Our findings point to substantial gene expression differences in lineage states between differentiated hCE and Caco-2 monolayers. Human small intestinal organoids and monolayers have been used to model TcdA/B toxicities; thus, we also compared barrier functions of hCE and hSIE monolayers. In prior work, we established a method for culturing healthy human jejunal monolayers and tracking cell maturation using transepithelial electrical resistance (TEER),^36^ a non-invasive quantitative technique that is widely accepted to measure epithelial integrity.^49^ Here, we applied similar principles to differentiated hCE (DC), hSIE, and Caco-2 monolayers and characterized the variation in barrier properties as measured by TEER between multiple donors **(Fig. 4A)**.

**Figure 4.**
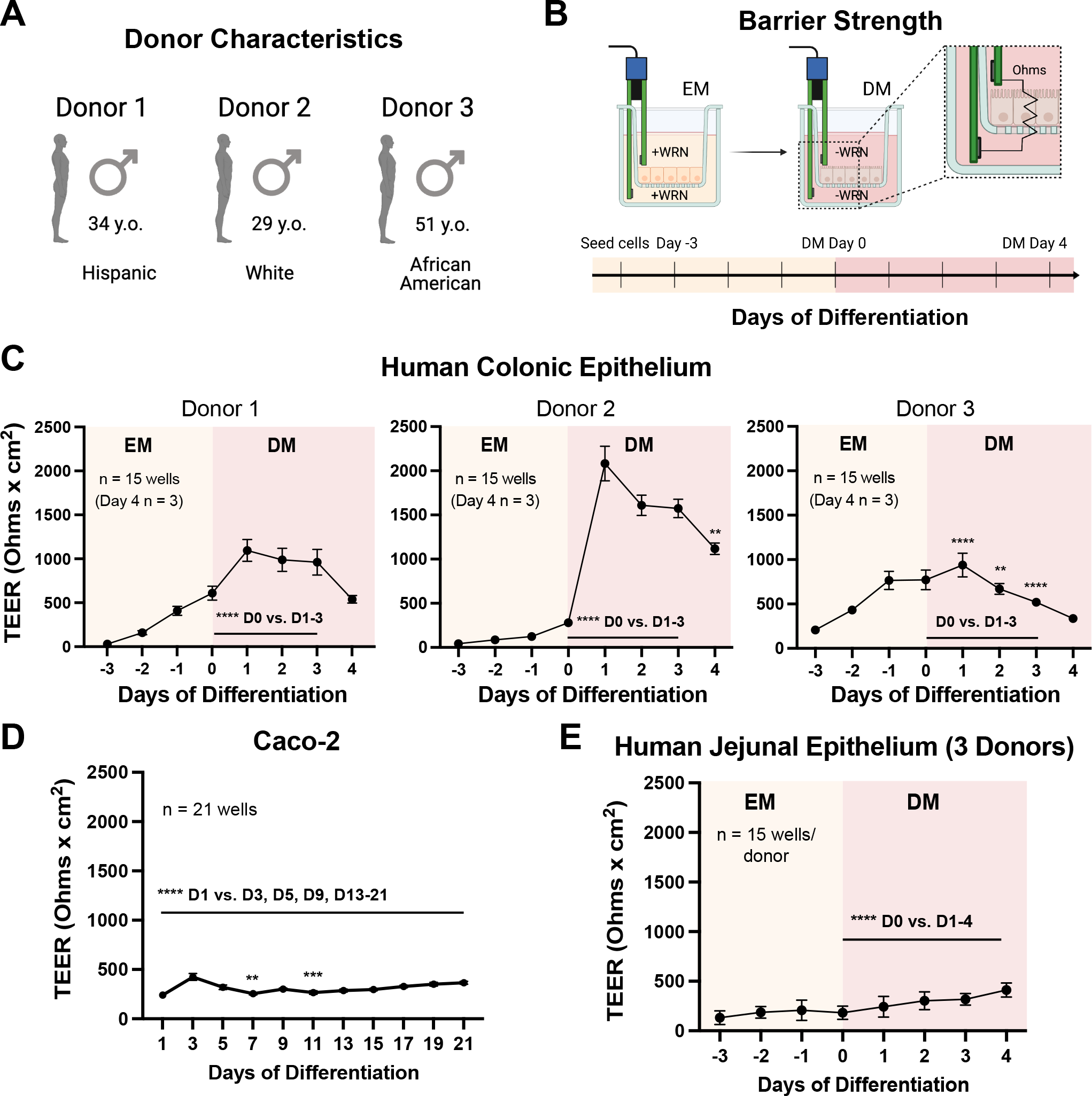
Human colonic epithelial cells exhibit elevated TEER during differentiation compared to Caco-2 or small intestinal epithelial monolayers. **(A)** Healthy human organ donor demographic information for transepithelial electrical resistance (TEER) assays. **(B)** Schematic and timeline for TEER (i.e., ionic conductance) measurements during differentiation of human intestinal epithelial monolayers. Monolayer TEER measurements for **(C)** human colonic epithelium, achieving a tight epithelial barrier within 1-day differentiation media (DM) exposure (N = 3 donors, n = 15 Transwells/donor for all days except n = 3 Transwells/donor for Day 4 DM), **(D)** Caco-2 cells (n = 21 Transwells), and **(E)** human small intestinal epithelium (N = 3 donors, n = 15 Transwells/donor). TEER values were compared against TEER at DM D0 (or Caco-2 Day 1) using a 1-way repeated measures ANOVA with Bonferroni correction. Each data point represents the mean ± SD. \**p* < 0.05, \*\**p* < 0.01, \*\*\**p* < 0.001, \*\*\*\**p* < 0.0001.

Daily TEER measurements were performed during monolayer expansion and differentiation **(Fig. 4B)**. In hCE monolayers from three healthy human donors, TEER was measured for 4 days in EM followed by 4 days in DM **(Fig. 4C)**. After just one day of DM exposure, hCE TEER reached an average value of 1000-2000 Ω × cm^2^, consistent with a tight epithelial barrier.^49^ While the individual TEER profiles varied by donor, the steep increase in hCE TEER after switching to DM was consistent across all three donors. It is noteworthy that both Caco-2 and hSIE monolayers did not reach high resistance like hCE cultures, with values staying below 500 Ω × cm^2^ **(Fig. 4D, E)**. This finding is consistent with previous jejunal TEER studies even after 12 days of differentiation,^36^ supporting the notion that hCE monolayers have a different barrier strength profile than hSIE or Caco-2 monolayers, which exhibit lower barrier strength profiles. Thus, differentiated Caco-2 cells, which have a baseline TEER that is much lower than that of differentiated hCE and is more consistent with ISCs and hSIE, may not accurately report *C. difficile* toxin sensitivities in the colon.

### Colonic Monolayers Exhibit a Dose-Dependent Early Cytotoxicity to TcdA, but Not TcdB, After Apical Exposure

By Days 2-4 of differentiation, hCE show transcriptional signs of maturation **(Fig. 3B, D)** and TEER is sufficiently elevated, indicating a tight barrier **(Fig. 4C)**. A drop in TEER at later timepoints is inherent to these differentiated primary culture systems since these cells undergo a natural ~5-day lifespan after leaving the stem/progenitor cell state. Interestingly, this may be an intrinsically programmed lifespan as mouse and human epithelial cells in vivo exhibit a similar lifespan.^50^ Given these features of the culture system, toxin experiments were initiated on Day 3 of differentiation at time t = 0 **(Fig. 5A)** as this was between 2 timepoints that closely mimic the in vivo maturation state, exhibits a relatively high barrier function as measured by TEER, and provides a sufficient window to perform toxicity assays prior to the cells’ terminal lifespan.

**Figure 5.**
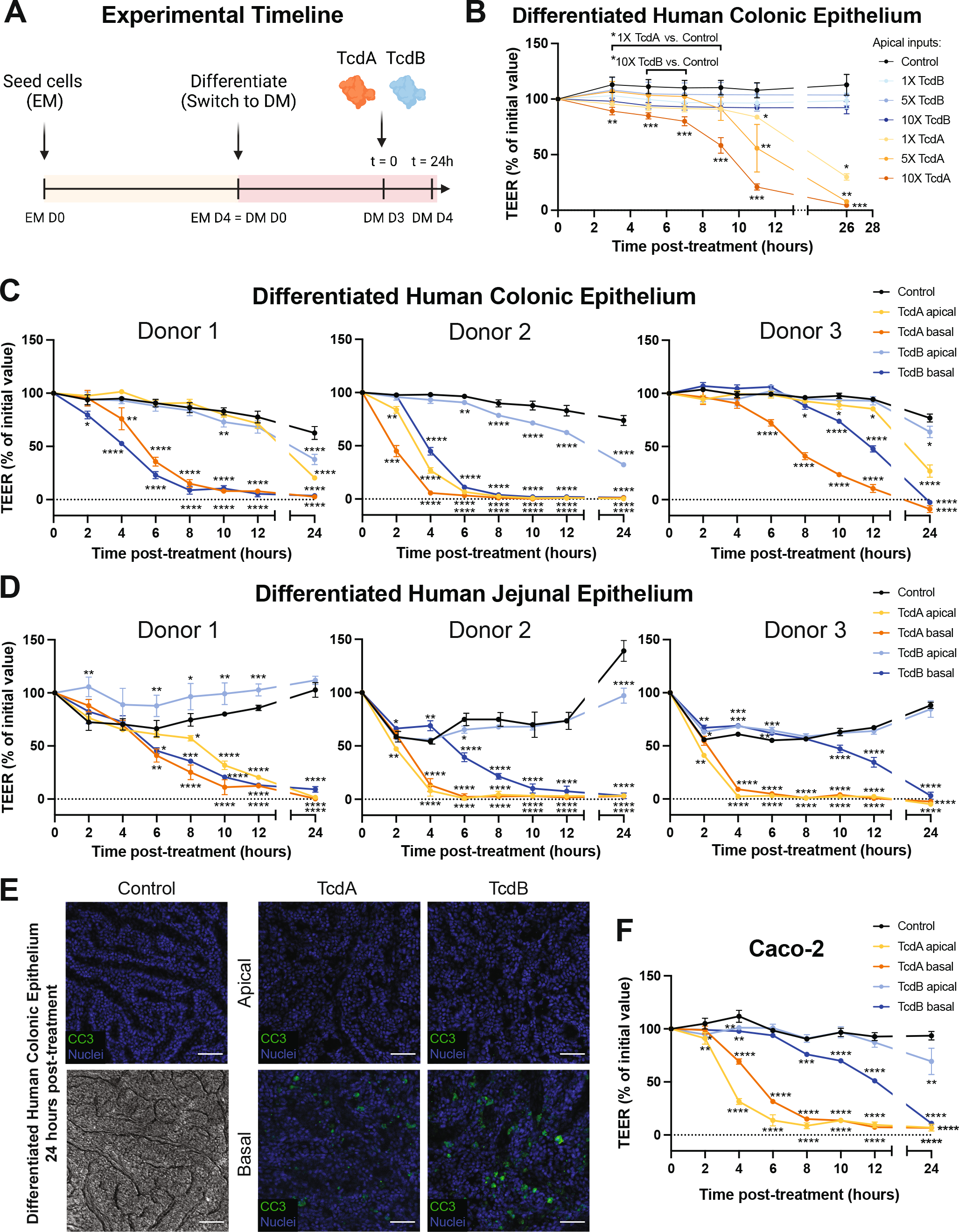
Basal *C. difficile* TcdA/B exposure induces more rapid cytotoxicity than apical toxin exposure in differentiated human colonic epithelium. **(A)** After seeding cells on Transwells in expansion media (EM) and switching to differentiation media (DM), TcdA or TcdB was added on DM Day 3 (D3) at t = 0 for 24 hours. **(B)** Dose-response of differentiated human colonic epithelium (hCE) to apical TcdA or TcdB at 30 pM (1X), 150 pM (5X), or 300 pM (10X); n = 4 Transwells, mean ± SD. TEER is reported as % of initial value (at t = 0). Kruskal-Wallis test with Benjamini-Hochberg correction was used. \**q* < 0.05, \*\**q* < 0.01, \*\*\**q* < 0.001. Monolayer TEER measurements for differentiated **(C)** hCE and **(D)** human jejunal epithelium. **(E)** Representative immunofluorescence staining (CC3, cleaved-caspase 3; Nuclei, bisbenzimide) and brightfield image (Control) of fixed apical/basal TcdA or TcdB-treated hCE, 24 hours post-treatment. Images were taken using a Zeiss LSM710 confocal 10X objective and ZEN 2011 acquisition software, with post-acquisition processing using FIJI. All scalebars = 50 μm. **(F)** TEER measurements for differentiated Caco-2 monolayers after apical or basal addition of TcdA or TcdB. For **C, D, and F**, toxins were added to DM at 1X = 30 pM; N = 3 donors for human colon and jejunum, and n = 3 Transwells, mean ± SD. 1-way ANOVA followed by Dunnett’s multiple comparisons test were used. \**p* < 0.05, \*\**p* < 0.01, \*\*\**p* < 0.001, \*\*\*\**p* < 0.0001.

To evaluate the sensitivity of the system to detect cytotoxic effects by TEER, three doses of TcdA and TcdB were applied apically to mature hCE monolayers. A dose of 30 pM was used as 1X as this is a physiological concentration identified in patients with active CDI.^51^ TcdA or TcdB at doses of 30 pM (1X), 150 pM (5X), and 300 pM (10X) were applied to differentiated hCE, and TEER changes were monitored during the first 12 hours and after 24 hours based on prior evidence of cytotoxicity within that timeframe **(Fig. 5B)**.^25,52^ hCE monolayers apically exposed to TcdB at any dose did not show a significant drop in TEER over 24 hours of exposure. Unlike TcdB, TcdA demonstrated a dose-dependent toxicity whereby 5X and 10X increasingly hastened the onset of barrier loss as measured by TEER **(Fig. 5B)**. These findings indicate that the hCE culture system is sufficiently sensitive to detect the impact of clinically relevant concentrations of TcdA and shows that TcdA apical exposure is more toxic than TcdB apical exposure at a molar-to-molar ratio.

### Basal Exposure to *C. difficile* Toxins A/B Produces More Rapid and Severe Cytotoxicity Than Apical Toxin Exposure

The lack of overt TcdB cytotoxicity when applied to the apical aspect of hCE monolayers was surprising given that TcdB is associated with severe toxicity in other systems.^22, 23, 53^ The observation that differentiated hCE monolayers were refractory to apical TcdB exposure prompted the question as to whether hCE and hSIE monolayers possess inherently different apical and basal toxin sensitivities. Etiologically, this is an important distinction to evaluate since our data imply that a healthy hCE monolayer may be highly resistant to apical *C. difficile* toxins. If TcdA/B have more potent effects when in contact with the basal aspect of cells, then CDI may primarily promote clinical sequelae when the epithelial barrier is compromised due to conditions that cause a leaky gut epithelium, at which point TcdA/B can access the basal aspect of the cell and cause rapid onset of toxicity. To evaluate differential sensitivities of apical and basal *C. difficile* toxin at a molar-to-molar ratio, 1X (30 pM) toxin was applied to either the apical or basal reservoirs of Transwells with differentiated hCE or hSIE monolayers from three separate human donors **(Fig. 5C, D)**.

TcdB applied to the apical aspect of differentiated hCE monolayers from three different donors demonstrated either no cytotoxicity or limited and delayed toxicity at 12 hours post-TcdB exposure **(Fig. 5C)**. By contrast, there was rapid and substantial cytotoxicity as early as 2 hours (on average, ~4.7 hours) when TcdB was applied to the basal aspect of the monolayers across all donors. Similarly, there was a trend of significantly delayed toxicity for TcdA when applied to the apical aspect of the monolayers compared to when TcdA was applied to the basal aspect of cells. For 2 donors, the monolayers demonstrated limited apical TcdA cytotoxicity at 12 hours, whereas basal exposure to TcdA resulted in rapid onset of cytotoxicity as early as 2 hours (on average, 4 hours) post-exposure across all three donors **(Fig. 5C)**. Interestingly, Donor 2 monolayers were more sensitive to apical application of both TcdA/B, and Donor 3 monolayers demonstrated delayed onset of cytotoxicity for both TcdA/B compared to the other two donor monolayers. Overall, TcdA/B demonstrated more cytotoxicity when exposed to the basal aspect of cells. Consistent with this, hCE monolayers were refractory to apoptosis at 24 hours after apical TcdA or TcdB exposure, while basal toxin exposure caused obvious signs of apoptosis **(Fig. 5E)**. Differential toxin sensitivities observed between donors is interesting and may reflect genetic backgrounds that render the differentiated hCE monolayers more or less responsive to the mechanisms of toxicity, which could be useful for determining patient-specific responses.

Single-cell RNAseq data of the *C. difficile* receptor gene expression profiles demonstrated that 2 out of the 3 known TcdA receptor genes (*SI* and *LRP1*) have significantly higher gene expression in mature absorptive lineages of the small intestine versus colon **(Fig. 2B)**. Thus, we hypothesized that differentiated hSIE monolayers would be more sensitive than differentiated hCE monolayers to TcdA. When TcdA was applied to the apical aspect of monolayers, substantial cytotoxicity was observed in hSIE monolayers as early as 2 hours (on average, 4 hours) **(Fig. 5D)**. By comparison, differentiated hCE demonstrated limited apical TcdA cytotoxicity in 2 donors with a late onset of 12-24 hours (on average, 12 hours across all donors) **(Fig. 5C)**. These findings show that differentiated hSIE is more sensitive to apical exposure to TcdA compared to differentiated hCE. Of note, TcdA/B cytotoxicity in Caco-2 monolayers is more consistent with trends observed in the hSIE monolayers compared to the hCE monolayers, with rapid sensitivity to apical TcdA within 2-4 hours and a relatively slower response to basal TcdB **(Fig. 5F)**. This highlights that, despite being derived from colon epithelial cancer, Caco-2 cells respond to toxins more like hSIE than hCE monolayers.

### A Leaky hCE Induced by NSAIDs Reduces Apical Barrier Protection From TcdB

Our results so far suggest limited sensitivity to TcdB on the apical/luminal surface of the human colon, but enhanced TcdB toxicity when applied to the basal surface. We hypothesized that toxin translocation from the apical to the basal side may be initiated or enhanced by a “leaky gut”, a term used to describe a broad range of underlying genetic, disease, or injury states that create increased intestinal paracellular permeability.^54^ Nonsteroidal anti-inflammatory drugs (NSAIDs) in general are associated with colitis,^55^ and diclofenac (DCF) in particular has demonstrated a significantly increased risk of *C. difficile*-associated diarrhea.^56,57^ Diclofenac, a commonly prescribed NSAID, has been previously shown to induce a leaky gut in primary hSIE.^58^ Clinically-relevant dosing of 1 mM was determined based on translating a typical therapeutic dose of 50 mg oral diclofenac into local concentrations of diclofenac between 300-1600 μM within the intestinal lumen.^59–61^

While TEER is an effective method to monitor ion flux across a damaged epithelial monolayer, it may not accurately report flux of toxin-sized molecules. To test whether a leaky gut induced by diclofenac initiates apical-to-basal translocation of toxin-sized molecules, a permeability assay was designed using hCE monolayers to monitor the translocation of conjugated dextrans from the apical to basal reservoir after toxin exposure **(Fig. 6)**. Large molecular weight dextrans should remain in the apical reservoir of the Transwells if there is a healthy epithelial barrier. FITC-Dextran (250 kDa) was chosen because it is similar to the molecular weight of TcdB (~270 kDa), which would provide insight into the time course in which a TcdB-sized particle would be able to pass paracellularly when the epithelial barrier is damaged. In the first 12 hours, apical diclofenac exposure alone induced modest leakiness which was compounded by the apical addition of TcdB, as measured by FITC-Dextran permeability **(Fig. 6)**. Of all the TcdB experimental conditions, increased permeability was only observed in the diclofenac/TcdB condition at 12 hours, consistent with the idea that epithelial barrier damage was allowing TcdB to interact with the basal aspect of the monolayer. At 24 hours, apical diclofenac/TcdB continued to show significantly increased permeability of FITC-Dextran, trending toward permeability levels that are only induced by direct application of TcdB to the basal side of the hCE monolayer **(Fig. 6)**. Taken together, these findings further reveal that a leaky colonic epithelial barrier can enhance the cytotoxic effects of apical TcdB exposure, likely through a mechanism whereby TcdB gains access to the basal aspect of the epithelial monolayer.

**Figure 6.**
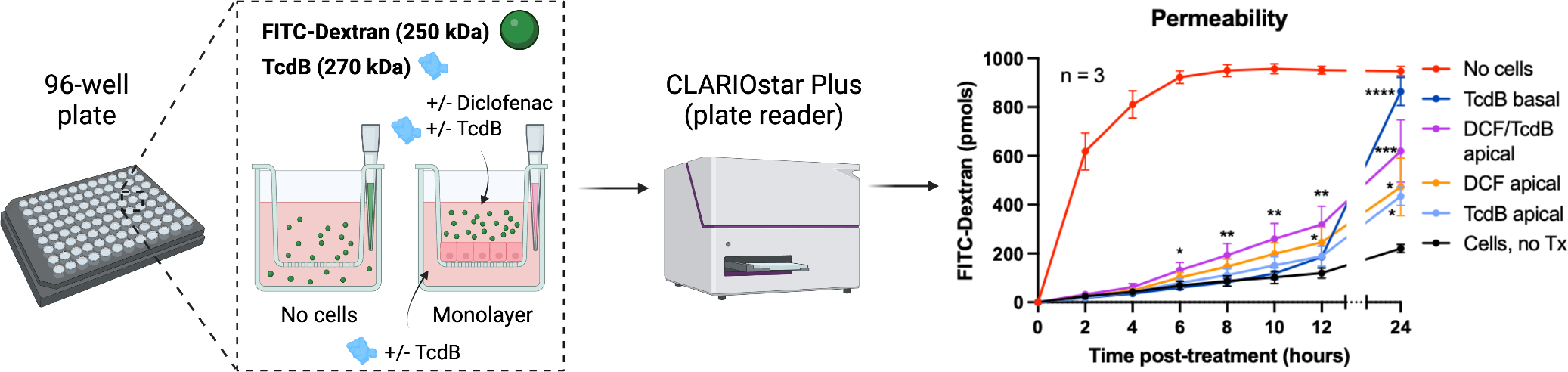
A leaky human colonic epithelium induced by diclofenac reduces apical barrier protection from TcdB. Schematic showing TcdB-sized FITC-Dextran (250 kDa) added to the apical surface of Transwells with or without differentiated human colonic epithelium (hCE), +/− TcdB, and +/− diclofenac and resulting permeability (n = 3 Transwells). Permeability data shows cumulative FITC-Dextran (pmol) transported from apical to basal compartments of Transwells. Tx = treatment. Each data point represents the mean ± SD. 1-way ANOVA followed by Dunnett’s multiple comparisons test were used. \**p* < 0.05, \*\**p* < 0.01, \*\*\**p* < 0.001, \*\*\*\**p* < 0.0001.

## DISCUSSION

In this work, we describe uncoupled apical and basal toxicity of *C. difficile* TcdA/B on hCE in multiple healthy human donors. We demonstrate that toxin sensitivities differ between hCE and hSIE, which could be related to different magnitudes of toxin receptor distribution across anatomical regions of the gut as revealed by our transcriptomic data. This interpretation is supported by findings that some toxin receptors are preferentially located on the apical versus basal surface of small intestinal or colonic epithelium.^28–33, 40^ The importance of identifying uncoupled apical/basal magnitude and time course of TcdA/B cytotoxicities may help inform the approach to develop more effective therapeutic modalities. As basal toxin exposure has more substantial cytotoxic effects, the use of luminal therapeutics such as probiotics or toxin-neutralizing drugs could be used as prophylaxis to prevent toxins from reaching the basal surface where they have more potent effects. This also has important clinical implications for patients with underlying intestinal epithelial barrier dysfunction, suggesting that they may have greater sensitivity to *C. difficile* toxins due to basal receptor interactions.

While 3D organoids and 2D monolayers are becoming more widely used to study host-microbe interactions, here we specifically tailor and characterize a first-in-kind 2D monolayer system in vitro that closely mimics the transcriptomic state of late ACCs found in vivo. We then focus the assay development in the system to detect the first signs of barrier loss caused by *C. difficile* toxins. Although preclinical intestinal models traditionally focus on genetically homogenous Caco-2 cells and other cell lines,16-19, 62-65 our transcriptomic data and barrier strength readouts show that Caco-2 cells remain in a relatively undifferentiated state and have a more similar barrier function profile to hSIE monolayers. This suggests that the hCE monolayers developed in our system are likely a more accurate preclinical model to evaluate toxin effects that are observed in the primary anatomical location in patients affected by CDI.^27^

The use of cancer cell lines such as Caco-2 as models to investigate biological mechanisms or preclinical models for drug testing comes with some advantages, such as ease/low cost of use and reproducible readouts. On the other hand, they suffer from homogenous responses that likely will not report the variability of responses observed in the human population. We view the inherent variability observed between hCE monolayers derived from different donors as an advantage as it has the potential to capture variability that exists in the human population. In this study, healthy transplant-grade tissue from three adult male donors varying in age and race/ethnicity was used. However, the ease of the hCE toxicity assay renders it scalable to additional donor demographics for increased representation of variable responses between humans. For example, Donor 2 appeared to be unusually sensitive to apically applied TcdA. While we cannot attribute a cause, unlike the other two donors, Donor 2 is White, and there is evidence to suggest that White patients demonstrate a higher CDI rate than patients of other races or ethnicities.^66^ While we acknowledge that sample sizes would have to be increased to confirm a statistical association, this example shows that our system allows sufficiently sensitive detection of toxin responses between humans. On-going efforts to bank ISCs from female and pediatric donors will allow us to evaluate the role of sex and age on *C. difficile* toxicity. Notably, this could increase the predictive value of our system as a preclinical model and address a substantial limitation of the genetically homogenous Caco-2 cell line.

Another major strength of this model is that it allows us to consider *C. difficile* toxin effects as a multi-step process: early barrier dysfunction (initial TEER drop) which leads to increased paracellular permeability (FITC-Dextran basal accumulation), cell rounding, and then eventual cytotoxic cell death (apoptosis) **(Fig. 7)**. Whereas currently used cell culture cytotoxicity assays that characterize cell rounding are subjective, non-standardized, and require 24 to 48 hours of incubation,^67^ our platform sensitively and quantitatively captures deleterious effects of TcdA/B in as little as 2 hours. Notably, while this platform was designed to evaluate the earliest events in toxicity on differentiated epithelium, the transcriptomic data show that this platform could also be used to evaluate the impact of toxins on the stem/undifferentiated cell lineages, which to-date has not been fully characterized but has important implications for wound healing and epithelial regeneration in CDI.^68^ Overall, this platform represents a flexible system and a strong experimental foundation to evaluate mechanisms of cell death or other aspects of cellular toxicities for development of new therapeutic approaches.

**Figure 7.**
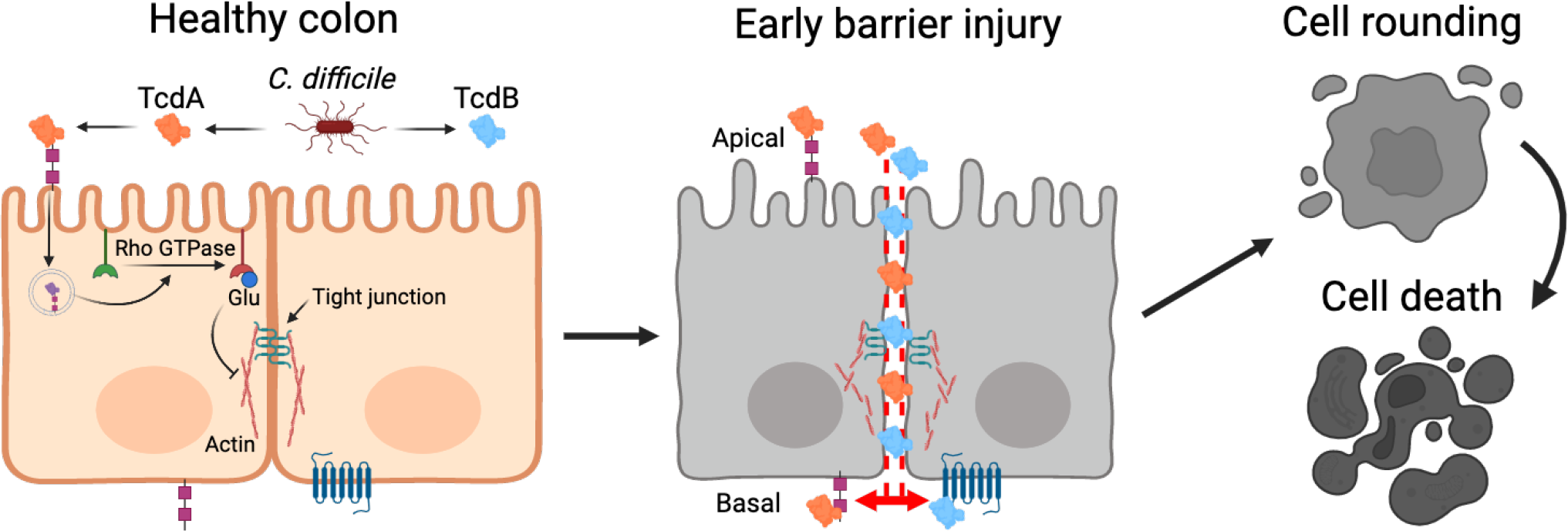
Human colonic epithelial cell platform detects early evidence of *C. difficile* toxicity. TcdA and TcdB enzymatic toxins are secreted by *C. difficile*, bind to host receptors, enter cells via endocytosis, and inactivate Rho-family GTPases by monoglucosylation. This results in early barrier injury, increased paracellular permeability to toxins, cell rounding, and eventual apoptosis.

Beyond studying *C. difficile* toxicity, the hCE platform established here can be utilized for a wide range of translational research applications, such as evaluating the toxicity of other pathogens, modeling a leaky/inflamed hCE, and drug screening. While we tailored the hCE monolayers in this study to detect and monitor the impacts of *C. difficile* toxins A/B, our methodological approach could be applied to other microbes. For example, *Vibrio cholerae* and toxigenic *Escherichia coli*, which both are associated with life-threatening diarrhea, produce toxins that impact other colonic physiological properties such as water and ion transport that could be further characterized using the transcriptomic and experimental tools highlighted in this study.^69,70^ Additionally, given that intestinal barrier defects are associated with a variety of human pathologies and diseases, the leaky gut model could be expanded to study other disease states associated with intestinal barrier defects including IBD, celiac disease, and irritable bowel syndrome^54^ to understand how these underlying conditions could potentiate disease etiology and negative effects of pathogenic microbes.

## METHODS

### Donor Selection

Human transplant-grade donor intestines were obtained from HonorBridge (Durham, NC) and exempted from human subjects research by the UNC Office of Human Research Ethics. Donor acceptance criteria were as follows: age 65 years or younger, brain-dead only, negative for human immunodeficiency virus, hepatitis, syphilis, tuberculosis, or COVID-19, as well as no prior history of severe abdominal injury, bowel surgery, cancer, or chemotherapy. Pancreas transplant donors were also excluded due to excision of proximal small intestinal tissue. Human intestinal tissue from three male donors (aged 29, 45, and 53 years) was used for scRNAseq.^39^ Colonic tissue from a 34-year-old Hispanic male was used for bulk RNAseq, immunofluorescence staining, and monolayer studies involving FITC-conjugated dextrans. Tissue from the same donor, in addition to tissue from a 51-year-old African American male and a 29-year-old Caucasian male, was used for TcdA and TcdB apical versus basal cell surface studies comparing colon and small intestine.

### Single-cell RNA Sequencing Processing and Analysis

Single-cell data was obtained from our previously published human intestinal dataset (Gene Expression Omnibus accession number: GSE185224), sequenced using an Illumina NextSeq 500 (Illumina, San Diego, CA).^39^ All processing and analyses were completed using scanpy (v1.7.2).^71^ Read counts were normalized to the median read depth of the data set and log-transformed.

### Bulk RNA Sequencing Preparation, Processing, and Analysis

To examine the changes in gene expression as colon stem cells differentiate into late absorptive colonocytes (ACCs) in vitro, RNAseq was performed on hCE monolayers that were harvested the day after seeding onto Transwells (EM Day 1), immediately prior to switching to DM (Day 0), and on Days 2 and 4 of differentiation on Transwells. For Caco-2, RNAseq was performed using cells harvested on Day 22 of differentiation. Three technical replicates were collected at each time point, and RNA was extracted via the RNAqueous-Micro Total RNA Isolation Kit (AM1931; ThermoFisher, Waltham, MA) according to the manufacturer’s protocol.

Before cDNA library preparation, RNA quality was determined by quantifying RNA integrity number (RIN), the ratio of 28s/18s RNA present, and RNA concentration in each sample. All samples used in library preparation had a RIN of at least 7. RIN was determined with the RNA 6000 Pico Kit for the Agilent 2100 Bioanalyzer. The Advanta RNA-Seq NGS Library Prep Kit for the Standard BioTools (formerly Fluidigm) Juno was used to prepare cDNA libraries for sequencing. Bulk sequencing was run on a NovaSeq 6000 instrument using one lane of an S4 v1.5 flow cell (Illumina), 2B paired-end reads, 150x read length. The Kallisto package was used to quantify transcript abundance which were pseudoaligned to human genome GRCh38. The output gene expression matrix was normalized using TMM normalization. Principal component analysis was performed with scikitlearn (v0.24.0, Rocquencourt, France).^72^ Gene set enrichment analysis (GSEA) was performed with normalized bulk RNAseq data using GSEA v4.2.3^73,74^ and previously published gene signatures for colon ISC and late ACC clusters versus the rest of the colon.^39^

### Collagen Hydrogel Scaffold Preparation

6-well tissue-culture plates (3516; Corning, Corning, NY) for expansion were coated with ice-cold type I rat tail collagen (354236; Corning, Corning, NY) in neutralization buffer based on an established protocol.^75^ Briefly, after being incubated at 37°C in 5% CO2 for 1 hour, sterile tissue-culture plates were coated with 1 mL of diluted (1 mg/mL) rat-tail collagen for each well. Plates were incubated at 37°C in 5% CO2 for 1 hour, then 3 mL room-temperature 1X Dulbecco’s phosphate-buffered saline (DPBS) (14040141; Thermo Fisher Scientific, Waltham, MA) was added onto each well. Plates were then kept overlaid with DPBS at room temperature for ≥ 2 weeks prior to use.

### Primary Human Crypt Isolation and Intestinal Epithelial Stem Cell Culture

Surgical specimens of human small and large intestines were obtained from donors at HonorBridge (formerly Carolina Donor Services, Durham, NC). Crypts from desired regions (Jej, DC) of each donor were detached from the specimen as previously described^36, 37, 39^ using a chelating buffer^76^ composed of EDTA (2 mM), dithiothreitol (DTT, 0.5 mM, freshly added), Na_2_ HPO_4_ (5.6 mM), KH_2_PO_4_ (8.0 mM), NaCl (96.2 mM), KCl (1.6 mM), sucrose (43.4 mM), D-sorbitol (54.9 mM), pH 7.4. Released crypts were expanded as a monolayer on a neutralized collagen hydrogel as described previously.^36–38^

Briefly, crypts were placed on the top of 1 mg/mL collagen hydrogels (1 ml into each well of 6-well plate) at a density of 10,000 crypts/well, overlaid with 3 mL of Expansion Media (EM) containing 10 mmol/L Y-27632 (S1049; SelleckChem, Houston, TX), and incubated at 37°C in 5% CO_2_. **See Table 1 for small intestine and colon EM formulations**. EM was used to expand the epithelial cell numbers as monolayers; media was changed the day after seeding and every 48 hours afterwards. When the cell coverage was greater than 80% (typically 4 to 6 days), the epithelium was dissociated to fragments to seed onto either 6-well tissue-culture plates coated with collagen hydrogels for continued expansion, or onto 12-well Transwell inserts (3460; Corning, Corning, NY) coated with 1% Matrigel for experiments.

**Table 1.**
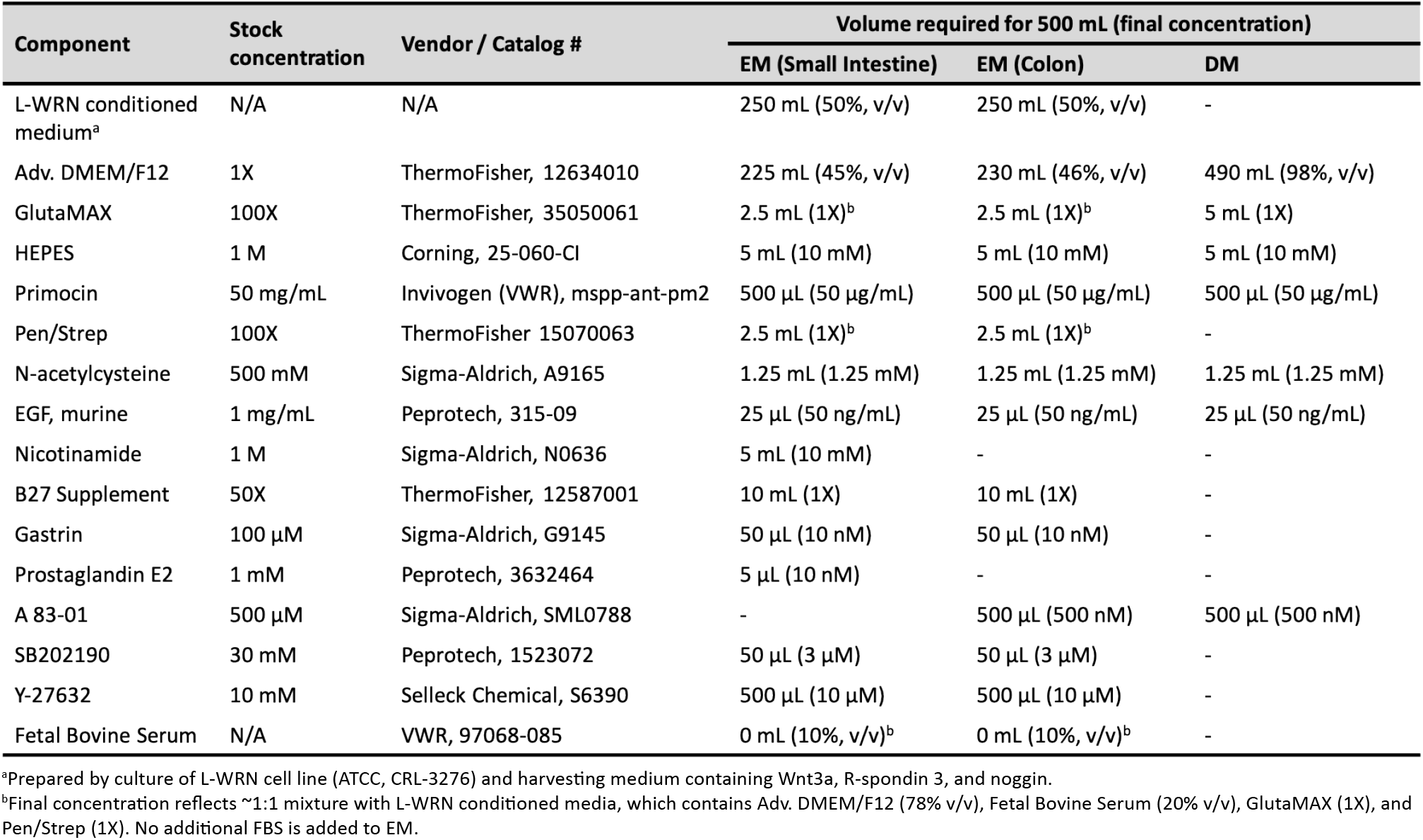
Formulations of primary cell culture media

| Component | Stock concentration | Vendor / Catalog # | Volume required for 500 mL (final concentration) |  |  |
| --- | --- | --- | --- | --- | --- |
|  |  |  | EM (Small Intestine) | EM (Colon) | DM |
| L-WRN conditioned medium<sup>a</sup> | N/A | N/A | 250 mL (50%, v/v) | 250 mL (50%, v/v) | - |
| Adv. DMEM/F12 | 1X | ThermoFisher, 12634010 | 225 mL (45%, v/v) | 230 mL (46%, v/v) | 490 mL (98%, v/v) |
| GlutaMAX | 100X | ThermoFisher, 35050061 | 2.5 mL (1X)<sup>b</sup> | 2.5 mL (1X)<sup>b</sup> | 5 mL (1X) |
| HEPES | 1 M | Corning, 25-060-Cl | 5 mL (10 mM) | 5 mL (10 mM) | 5 mL (10 mM) |
| Primocin | 50 mg/mL | Invivogen (VWR), msp-p-ant-pm2 | 500 µL (50 µg/mL) | 500 µL (50 µg/mL) | 500 µL (50 µg/mL) |
| Pen/Strep | 100X | ThermoFisher 15070063 | 2.5 mL (1X)<sup>b</sup> | 2.5 mL (1X)<sup>b</sup> | - |
| N-acetylcysteine | 500 mM | Sigma-Aldrich, A9165 | 1.25 mL (1.25 mM) | 1.25 mL (1.25 mM) | 1.25 mL (1.25 mM) |
| EGF, murine | 1 mg/mL | Peprotech, 315-09 | 25 µL (50 ng/mL) | 25 µL (50 ng/mL) | 25 µL (50 ng/mL) |
| Nicotinamide | 1 M | Sigma-Aldrich, N0636 | 5 mL (10 mM) | - | - |
| B27 Supplement | 50X | ThermoFisher, 12587001 | 10 mL (1X) | 10 mL (1X) | - |
| Gastrin | 100 µM | Sigma-Aldrich, G9145 | 50 µL (10 nM) | 50 µL (10 nM) | - |
| Prostaglandin E2 | 1 mM | Peprotech, 3632464 | 5 µL (10 nM) | - | - |
| A 83-01 | 500 µM | Sigma-Aldrich, SML0788 | - | 500 µL (500 nM) | 500 µL (500 nM) |
| SB202190 | 30 mM | Peprotech, 1523072 | 50 µL (3 µM) | 50 µL (3 µM) | - |
| Y-27632 | 10 mM | Selleck Chemical, S6390 | 500 µL (10 µM) | 500 µL (10 µM) | - |
| Fetal Bovine Serum | N/A | VWR, 97068-085 | 0 mL (10%, v/v)<sup>b</sup> | 0 mL (10%, v/v)<sup>b</sup> | - |

<sup>a</sup>Prepared by culture of L-WRN cell line (ATCC, CRL-3276) and harvesting medium containing Wnt3a, R-spondin 3, and noggin. <sup>b</sup>Final concentration reflects ~1:1 mixture with L-WRN conditioned media, which contains Adv. DMEM/F12 (78% v/v), Fetal Bovine Serum (20% v/v), GlutaMAX (1X), and Pen/Strep (1X). No additional FBS is added to EM.

### Transwell Preparation and Intestinal Epithelial Stem Cell Differentiation

Briefly, 500 μL ice-cold 1% Growth Factor Reduced Matrigel (354230; Corning, Corning, NY) diluted in ice-cold DPBS was added to the apical surface of 12-well Transwell inserts. Transwell plates were then incubated at 37°C in 5% CO_2_ overnight and then rinsed 2-3 times with 1X DPBS. Dissociated intestinal epithelial stem cells that were expanded on collagen hydrogels were resuspended in 500 μL of EM per Transwell insert and seeded on the apical surface of each insert at an approximate density of 1.5 × 10^5^ cells/cm^2^. 1.5 mL of EM was added to the basal reservoir at the time of seeding. Both the apical and basal media were replaced with fresh EM the day after seeding. After sufficient cell coverage on Transwells (e.g., no visible monolayer gaps), on Day 4, Differentiation Medium (DM)^77^ **(Table 1)** was used to initiate differentiation as well as during toxin experiments.

### Caco-2 Cell Culture and Differentiation

Caco-2 cells (HTB-37; ATCC, Manassas, VA) were grown on a 10 cm polystyrene tissue culture-treated dish (430167; Corning, Corning, NY) in Dulbecco’s Modified Eagle Media + 10% fetal bovine serum + 1% penicillin-streptomycin based on an established protocol **(Table 2)**.^78^ Caco-2 media was changed every 2 days until desired confluency was reached. Two 60% confluent 10 cm dishes were seeded onto 24 inserts in a 12-well Transwell plate coated with 30 μg/mL collagen in 1X DPBS. Spontaneous differentiation of Caco-2 monolayers was allowed to occur over 21 days, and media was changed every 2 days until the start of toxin experiments on Day 22.

**Table 2.** Formulation of Caco-2 media

| Component | Stock concentration | Vendor / Catalog # | Volume required for 500 mL (final concentration) |
| --- | --- | --- | --- |
| DMEM | 1X | ThermoFisher, 11995065 | 445 mL (89%, v/v) |
| Fetal Bovine Serum | N/A | VWR, 97068-085 | 50 mL (10%, v/v) |
| Pen/Strep | 100X | ThermoFisher, 15070063 | 5 mL (1X) |

### *C. difficile* Toxins Information

Toxin A (SML1154; Sigma-Aldrich, St. Louis, MO) and Toxin B (SML1153; Sigma-Aldrich, St. Louis, MO) were reconstituted in purified water and then diluted to 30 pM, 150 pM, or 300 pM (for dose-response) in DM before adding to differentiated cells or Caco-2 on Transwell plates. These native toxins are purified from *C. difficile* strain VPI10463 (toxinotype 0) which is from Clade 1 (as opposed to Clade 2 strains for which TcdB does not appear to bind FZDs).^79^ 30 pM (1X) was chosen as a clinically-relevant concentration based on ultrasensitive digital enzyme-linked immunosorbent assays (ELISAs) for TcdA and TcdB conducted on clinical specimens that tested positive for cytotoxicity.^51^

### Transepithelial Electrical Resistance (TEER) Measurements

Intestinal epithelial barrier integrity was quantified via transepithelial electrical resistance (TEER) using an EVOM2™ paired with STX2 electrodes (World Precision Instruments, FL). TEER was measured daily during differentiation and every 2 hours after adding toxin for 12 hours, followed by a 24-hour timepoint. Toxin was added on DM Day 3 (DC) or DM Day 4 (jejunum). For Caco-2, TEER was measured every two days (starting day 1) and media was changed every 2 days during spontaneous differentiation. For Caco-2 TEER measurements in the presence of toxin, TcdA or TcdB was added to the apical side of Transwell inserts on Day 22.

### Microscopy and Immunofluorescence Staining of Differentiated Human Colonic Epithelium

After the 24-hour timepoint, control and TcdA or TcdB-treated primary human colonic epithelial monolayers were fixed in 4% paraformaldehyde in PBS at 37°C for 15 minutes. Cells were then permeabilized in 0.5% Triton-X100 at room temperature for 20 minutes, followed by blocking with 3% BSA in PBS for 1 hour at room temperature. The cells were treated with primary antibody against cleaved-caspase 3 (1:300, 9661S; Cell Signaling Technology, Danvers, MA) and incubated at 4°C overnight. The cells were then washed with a rinse solution consisting of 0.1% BSA, 0.2% Triton-X100, and 0.05% Tween-20, then incubated with Alexa Fluor 488-conjugated donkey anti-rabbit antibody (1:500, 711-545-152; Jackson ImmunoResearch, West Grove, PA) and bisbenzimide H 33258 (1:1000, B1155; Sigma-Aldrich-Aldrich, St. Louis, MO) diluted in 3% BSA in PBS for 1 hour at room temperature. Cells were washed in 1X DPBS twice prior to microscopy. Images were captured using an LSM700 confocal microscope (Zeiss, Jena, Germany) 10X objective and ZEN 2011 acquisition software, with post-acquisition processing using FIJI.^80^

### Basal Fluorescent Dextran Quantification and Diclofenac Permeability Assays

Permeability assays were conducted as previously described.^36^ Briefly, 250 kDa FITC-Dextran (FD250S; Sigma-Aldrich, St. Louis, MO) was chosen to mimic the size of TcdB (270 kDa). FITC-Dextran was diluted to 1 μM in DM and added to the apical side of 12-well Transwell inserts with or without 30 pM TcdB. Basal media (100 μL) was collected every 2 hours for 12 hours and replaced with an equivalent volume of DM (or DM with TcdB), followed by a 24-hour timepoint.

Basal collections were added to a 96-well microplate for fluorescence-based assays (M33089; ThermoFisher, Waltham, MA). FITC-associated fluorescence was measured by CLARIOstar Plus Microplate Reader (BMG Labtech, Ortenberg, Germany) after excitation at 483 nm and detection at 530 nm. Relative fluorescence units were converted to picomoles using a standard curve. Diclofenac sodium salt (157660; MP Biomedicals, Solon, OH) was reconstituted in DMSO and diluted to 1 mM in DM containing FITC-Dextran with or without TcdB.

### Statistics

For comparing gene expression in scRNAseq data **(Fig. 2B, D, and F)**, significance was calculated by negative binomial regression with the diffxpy python package (v0.7.4) using the Wald test and Benjamini-Hochberg correction. TEER values at each day of DM (or Caco-2 media) were compared against TEER at D0 (or Caco-2 Day 1) using a 1-way repeated measures ANOVA with Bonferroni correction **(Fig. 4C-E)**. Experimental TEER values were compared against vehicle control for dose-response data **(Fig. 5B)** using the Kruskal-Wallis test with Benjamini-Hochberg correction due to data non-normality, and in **Fig. 5C, D, and F** using a 1-way ANOVA followed by Dunnett’s multiple comparisons test. 1-way ANOVA followed by Dunnett’s multiple comparisons tests were used to compare FITC-Dextran permeability between control and experimental groups **(Fig. 6)**.

## Abbreviations used in this paper

ACC: absorptive colonocyte
AE: absorptive enterocyte
CDI: *Clostridioides difficile* infection
CDT: *Clostridioides difficile* transferase
DC: descending colon
DCF: diclofenac
DM: differentiation media
EM: expansion media
FAE: follicle-associated epithelium
hCE: human colonic epithelium
hSIE: human small intestinal epithelium
IBD: inflammatory bowel disease
ISC: intestinal stem cell
NSAID: nonsteroidal anti-inflammatory drug
PC: principal component
scRNAseq: single-cell RNA-sequencing
TA: transit-amplifying
TcdA: Toxin A
TcdB: Toxin B
TEER: transepithelial electrical resistance
3D: 3-dimensional

## Access to Data

All authors had access to all the data and have reviewed and approved the final manuscript.

## ACKNOWLEDGEMENTS

First and foremost, thank you to the anonymous organ donors, their families, and HonorBridge (Durham, NC) for providing the intestinal tissue samples used in this study. Schematics in Figures 1A, 3A, 4A, 4B, 5A, 6, and 7 were created using BioRender. Thank you to the UNC High-Throughput Sequencing Facility and the CGIBD Advanced Analytics Core, especially Gabrielle Cannon, for RNA-seq data. Thank you to Casey Theriot, Rita Tamayo, Carly Catella, Carol Hall, Sudeep Sarma, Nick Markham, and Aadra Bhatt for useful discussions and James Stroud, Nathan Kohn, and Yunan Hu for support with TEER measurements.

